# A highly-contiguous genome assembly of the inbred Babraham pig (*Sus scrofa*) quantifies breed homozygosity and illuminates porcine immunogenetic variation

**DOI:** 10.1101/2023.10.04.560872

**Authors:** John C. Schwartz, Colin P. Farrell, Graham Freimanis, Andrew K. Sewell, John A. Hammond, John D. Phillips

## Abstract

The inbred Babraham pig serves as a valuable biomedical model for research due to its high level of homozygosity, including in the major histocompatibility complex (MHC) loci and likely other important immune-related gene complexes, which are generally highly diverse in outbred populations. As the ability to control for this diversity using inbred organisms is of great utility, we sought to improve this resource by generating a long-read whole genome assembly of a Babraham pig. The Babraham genome was *de novo* assembled using PacBio long-reads and error-corrected using Illumina short-reads. The assembled contigs were then mapped to the current porcine reference assembly, Sscrofa11.1, to generate chromosome-level scaffolds. The resulting Babraham pig assembly is nearly as contiguous as Sscrofa11.1 with a contig N50 of 34.95 Mb and contig L50 of 23. The remaining sequence gaps are generally the result of poor assembly across large and highly repetitive regions such as the centromeres and tandemly duplicated gene families, including immune-related gene complexes, that often vary in gene content between haplotypes. We also further confirm homozygosity across the Babraham pig MHC and characterize the allele content across several immune-related gene complexes, including the contiguous assemblies of the antibody heavy chain locus and leukocyte receptor complex. The Babraham pig genome assembly provides an alternate highly contiguous porcine genome assembly as a resource for the livestock genomics community. The assembly will also aid biomedical and veterinary research that utilizes this animal model such as when controlling for genetic variation is critical.

## Introduction

Pigs (*Sus scrofa*) are vital to both biomedical research and the production of pork, the most extensively consumed meat product worldwide (USDA 2022). The anatomical and physiological similarities with humans make pigs an excellent model of human disease, such as for tuberculosis or influenza (Bolin et al. 1997; Groenen et al. 2012; Perleberg et al. 2018; Holzer et al. 2021), and their similar organ sizes make pigs ideally suited as a source of organs for xenotransplatation (Lunney 2007; Ekser et al. 2017). Furthermore, pigs continue to face ongoing threats from African swine fever and other diseases, especially in east Asia, and research into effectively controlling these diseases is important for global food security and for improving animal welfare (Kedkovid et al. 2020).

The pig reference genome assembly (Groenen et al. 2012; Warr et al. 2020) has greatly contributed to our understanding of porcine immunology (Dawson et al. 2013; Schwartz et al. 2017; Massari et al. 2018; Morgan et al. 2018; Schwartz and Hammond 2018; Hammer et al. 2020; Zhang et al. 2020; Le Page et al. 2021; Linguiti et al. 2022) and has helped researchers better utilize the pig as a model of disease (Groenen et al. 2012; Nicholls et al. 2016; Perleberg et al. 2018). It has also facilitated the generation of genome-edited pigs, such as, for example, for resistance to porcine reproductive and respiratory syndrome virus (PRRSV) infection (Whitworth et al. 2014; Burkard et al. 2018), or for the inactivation of porcine endogenous retroviruses in order to improve the safety of xenotransplantation (Niu et al. 2017; Niu et al. 2021). Improvements in long-read sequencing technologies and whole genome assembly techniques within the last decade, however, have resulted in greatly improved mammalian genome assemblies, with contig lengths now approaching that of whole chromosomes, and at a greatly reduced financial cost (Bickhart et al. 2017; Koren et al. 2018; Low et al. 2020; Rice et al. 2020; Rosen et al. 2020; Warr et al. 2020; Bredemeyer et al. 2021). Among these endeavors, the pig reference genome was recently updated with Illumina paired-end reads, complete bacterial artificial chromosome (BAC) sequences, BAC and fosmid end sequences, and Pacific Biosciences (PacBio) single molecule real-time sequencing reads. While these sequences were generated using genomic DNA from the same purebred Duroc sow used for the earlier pig reference assembly, additional Y-chromosome sequence from another individual was incorporated into the current assembly, Sscrofa11.1 (Warr et al. 2020).

As an animal model with a defined genetic background and limited heterozygosity, the inbred Babraham pig holds great potential for the research community, and several recent studies have used it to investigate immune responses in the pig while leveraging the breed’s minimal genomic variability (Lefevre et al. 2012; Nicholls et al. 2012; Nicholls et al. 2016; Tungatt et al. 2018; Baratelli et al. 2020; Edmans et al. 2021; Martini et al. 2021). The breed was initially developed from commercial Large White pigs at The Babraham Institute (Cambridge, UK) in the 1970s as a model organism, and is currently the only extant large inbred pig breed available for research (Schwartz et al. 2018). Individuals were selectively bred to display the least amount of cross-rejection after multiple skin grafts, eventually producing animals with full cross-tolerance (Signer et al. 1999). Such graft tolerance suggested homozygosity across the major histocompatibility complex (MHC), which was later confirmed (Signer et al. 1999; Nicholls et al. 2016; Schwartz et al. 2018); and restriction fragment length polymorphism patterning also further indicated a level of inbreeding comparable to that of inbred mice (Signer et al. 1999).

Pigs are natural hosts of Influenza A virus (IAV) and infection represents a substantial problem for the agricultural industry (Brown 2000). Pigs can be infected with human and bird forms of IAV which can recombine with swine virus to generate antigenic shift and create dangerous pandemic strains (Ito et al. 1998; Ma et al. 2009). The Babraham pig has become an important model for understanding human influenza infection and for the development of new vaccines against IAV and other swine viruses (Lefevre et al. 2012; Rajao and Vincent 2015). The dominant influenza peptide antigens presented by Babraham MHC molecules (also known as swine leukocyte antigen (SLA)) have been described and peptide-SLA multimers have been used to study spatial, temporal, and molecular dynamics of swine flu-specific CD8+ tissue resident T-cells (Martini et al. 2022) and assess responses to IAV vaccines (Martini et al. 2021; Goatley et al. 2022). The absence of detailed architectural knowledge of the Babraham antigen receptor loci remains the major bottleneck in the Babraham model of viral infection. We set out to bridge this critical knowledge gap to bring this swine model to the level of understanding available in human or laboratory mice.

To improve the Babraham pig as a resource for transcriptomic and immunological studies, we utilized PacBio long-read sequencing and assembly, Illumina short-read error correction, and reference-guided scaffolding to generate a highly contiguous genome assembly of the inbred Babraham pig that is almost as contiguous as the reference assembly. To reduce the effect that somatically rearranging immune receptors might have on the assembly of the B cell and T cell receptor loci, we used brain tissue for whole genome sequencing and assembly due to the lack of lymphocytes generally present in that tissue. As immune-related gene complexes often contain many tandemly duplicated paralogous genes that can be highly similar in sequence and of variable gene content, their repetitiveness often disrupts genome assemblies (Bickhart et al. 2017; Rosen et al. 2020). We therefore specifically investigated the homozygosity, contiguity, and gene content of several highly variable regions that are important in lymphocyte immunobiology, including the B cell (IGH, IGK, and IGL) and T cell receptor (TRB, TRG, TRA/TRD) loci, the MHC class I and class II, the natural killer complex (NKC), and the leukocyte receptor complex (LRC) and compared them to previous characterizations in the pig.

## Results

### A highly contiguous *de novo* assembly of the Babraham pig genome

Approximately 1.11 x 10^7^ PacBio Sequel II reads with an average read length of 12,552 bp and read N50 of 22,299 bp were generated, amounting to approximately a 57-fold coverage of the porcine genome. Reads were *de novo* assembled into contigs and scaffolds using Flye (v2.5) (Kolmogorov et al. 2019) and error-corrected using Pilon (version 1.24) (Walker et al. 2014) and approximately 51-fold coverage of Illumina (2 x 150 bp) reads from the same animal. Contigs were then screened for contaminating sequence using Kraken (version 1.1.1) (Wood and Salzberg 2014). However, this did not identify any contamination and all contigs either successfully mapped to Sscrofa11.1 or contained simple repeats. The resulting assembly consists of 2,447 Mb across 1,391 contigs with a contig N50 of 34.95 Mb and contig L50 of 23. The assembled contigs and scaffolds were mapped to the pig reference genome assembly, Sscrofa11.1 (Warr et al. 2020) to generate a chromosome-level assembly (Table 1). This resulted in a placement of 357 contigs spanning 2,408 Mb across the 18 autosomes, Chr X, Chr Y, and the mitochondrial chromosome. The remaining 1,034 unplaced contigs, comprising 40 Mb, were generally much smaller with a contig N50 of 150 kb and are presumably mostly comprised of unplaced Chr Y sequence and alternative haplotype sequences.

**Table 1.** Chromosome-level assembly statistics for Sscrofa11.1, USMARCv1.0, and TPI_Babraham_pig_v1.

| Chr | Sscrofa11.1 |  |  | USMARCv1.0 |  |  | TPI Babraham pig_v1 |  |  |
| --- | --- | --- | --- | --- | --- | --- | --- | --- | --- |
|  | Ungapped length (bp) | Number of Contigs | Contig N50 (bp) | Ungapped length (bp) | Number of Contigs | Contig N50 (bp) | Ungapped length (bp) | Number of Contigs | Contig N50 (bp) |
| 1 | 274330132 | 5 | 90927422 | 268199312 | 66 | 6467034 | 278785404 | 15 | 60958165 |
| 2 | 151800670 | 7 | 87417173 | 141039314 | 37 | 8371877 | 150500149 | 53 | 36467456 |
| 3 | 132648513 | 9 | 73254198 | 128651370 | 35 | 6713922 | 133369682 | 10 | 21452984 |
| 4 | 130870669 | 4 | 100518328 | 128001252 | 30 | 11944967 | 131187407 | 12 | 34019294 |
| 5 | 104375107 | 13 | 21111347 | 98929882 | 27 | 6762634 | 107091043 | 11 | 17630615 |
| 6 | 170419461 | 11 | 18397423 | 160955110 | 41 | 7899165 | 170189100 | 18 | 20338316 |
| 7 | 121743199 | 12 | 29790190 | 119961677 | 18 | 29051871 | 125629992 | 26 | 22695760 |
| 8 | 138865937 | 6 | 72677949 | 135855389 | 33 | 6513734 | 139153575 | 15 | 76299722 |
| 9 | 139511883 | 3 | 133627600 | 135417841 | 27 | 10245400 | 139228276 | 15 | 41712811 |
| 10 | 69257333 | 4 | 44332889 | 68415272 | 22 | 5169423 | 70201652 | 9 | 10942133 |
| 11 | 79119678 | 5 | 19474953 | 77145484 | 16 | 6656154 | 79744139 | 5 | 29255778 |
| 12 | 61500128 | 4 | 45299297 | 56950340 | 19 | 4652640 | 61381331 | 12 | 18011077 |
| 13 | 208234567 | 11 | 24026255 | 199810805 | 50 | 7414089 | 208105327 | 14 | 48808734 |
| 14 | 141755246 | 3 | 130192676 | 139163928 | 16 | 38681723 | 141998002 | 15 | 34948847 |
| 15 | 140362525 | 4 | 38129723 | 136633314 | 30 | 7208255 | 140491208 | 15 | 81360748 |
| 16 | 79944280 | 1 | 79944280 | 77627177 | 24 | 5327307 | 80090520 | 6 | 44193870 |
| 17 | 63343681 | 8 | 48231277 | 62541674 | 14 | 8416923 | 63352208 | 14 | 48195611 |
| 18 | 55982971 | 1 | 55982971 | 55717653 | 7 | 30967442 | 55898280 | 2 | 48439141 |
| X | 125778901 | 11 | 16842758 | 99540920 | 159 | 960692 | 125650420 | 63 | 4139144 |
| Y | 17132043 | 413 | 66937 | 2610849 | 9 | 255686 | 5552120 | 26 | 635153 |
| M | 16613 | 1 | 16613 | 16760 | 1 | 16760 | 16701 | 1 | 16701 |
| Unplaced | 65054210 | 583 | 250081 | 329944915 | 14137 | 23975 | 39999133 | 1034 | 149592 |
| **TOTAL** | **2472047747** | **1119** | **48231277** | **2623130238** | **14818** | **6372407** | **2447615669** | **1391** | **34948847** |

The contiguity across the autosomes and Chr X are comparable between the Babraham and the Sscrofa11.1 assemblies (Figure 1). The allosomes, Chr X and Chr Y, are the least contiguous, and while the former is approximately the same length in the two assemblies, the Babraham Chr Y assembly is only 32 % the total length of Chr Y in Sscrofa11.1, indicating that a large proportion of Chr Y likely remains unplaced in the Babraham assembly. Fifty of the unplaced contigs mapped at least partially to the Chr Y assembly of Sscrofa11.1; however, the combined size of these contigs totaled only 2.4 Mb, indicating that a considerable amount of Chr Y remains unaccounted for. Sequence orientation and contig order were further confirmed for the autosomes and Chr X by mapping their assemblies back to Sscrofa11.1 (Figure 2).

**Figure 1.**
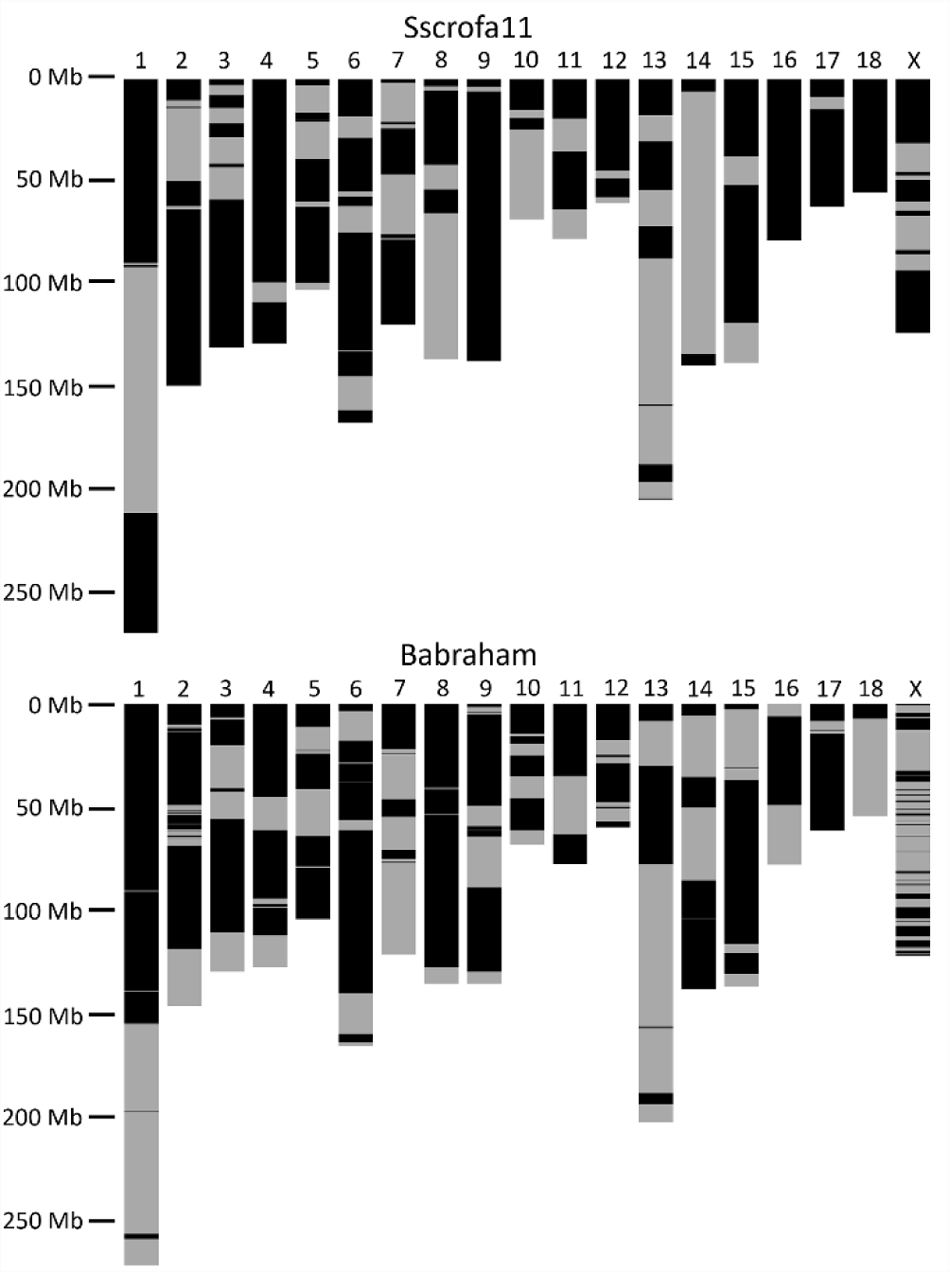
Contiguity of Sscrofa11.1 (*top*) and TPI_Babraham_pig_v1 (*bottom*) autosomal and Chr X assemblies. Contigs are indicated by alternating dark and light bands. Contigs smaller than 100 kb are not shown as they are too small to reasonably resolve.

**Figure 2.**
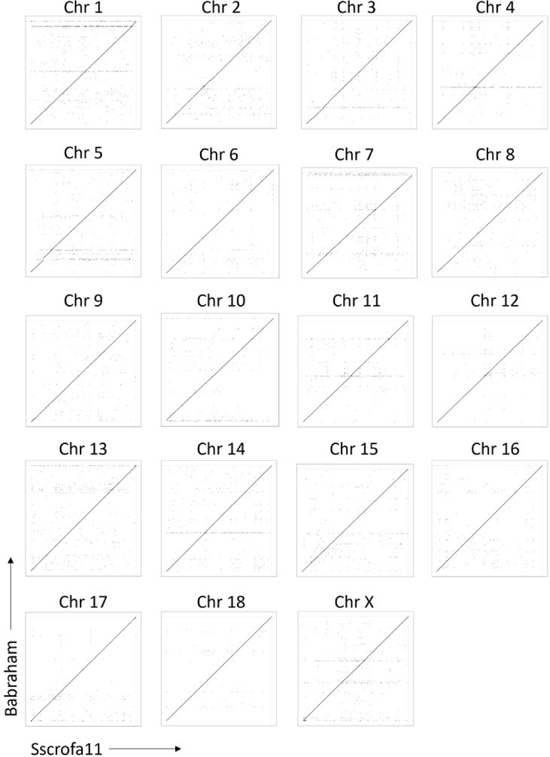
Recurrence plot comparisons of TPI_Babraham_pig_v1 (*vertical axes*) and Sscrofa11.1 (*horizontal axes*) autosomal and Chr X assemblies.

### Centromeric and telomeric repeats disrupt the sequence contiguity of the assembly

We next attempted to determine the degree of contiguity loss due to large and repetitive sequences, specifically the telomeres and centromeres, as these are likely to disrupt assembly contiguity. We detected centromeric repeats in the expected locations for all but three autosomes (Chr 10, Chr 12, and Chr 18) and Chr X, in which the centromeres were not identified (Figure 3). Of the remaining, all are disrupted by either a sequence gap (Chr 1 to Chr 12) or truncated, as is the case for the telocentric chromosomes (Chr 13 to Chr 18). Thus, not unexpectedly, the large repeat structures associated with the centromeres were problematic for the contiguous assembly of the genome. Furthermore, as noted for the Sscrofa11.1 assembly (Warr et al. 2020), the centromere of Chr 17 is found on the opposite end of the assembly as conventionally presented. However, to conform to the published reference assembly, we retained this reversed orientation for Chr 17.

**Figure 3.**
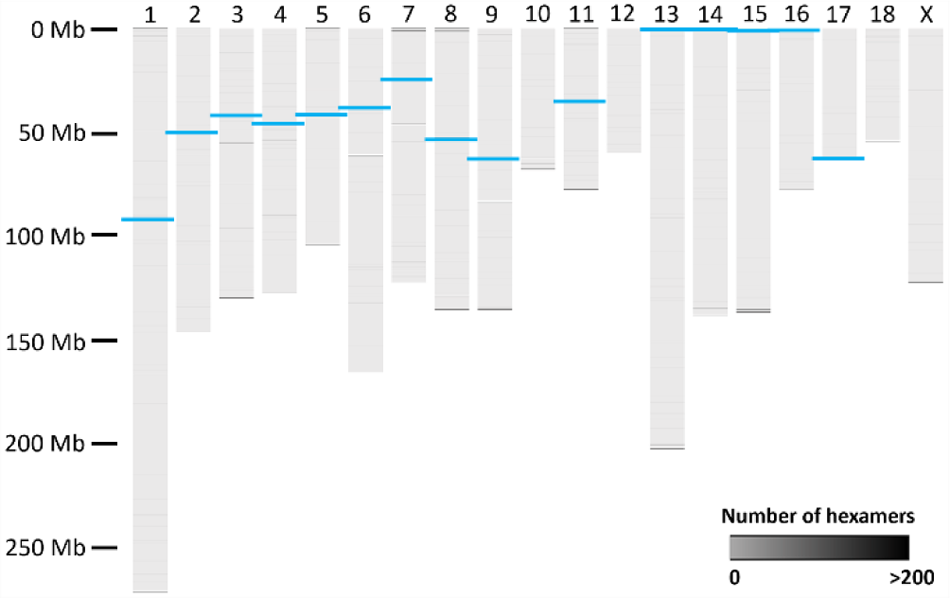
Centromeric and telomeric repeats in the TPI_Babraham_pig_v1 assembly. Repeats of telomeric hexamers are shown as *grey* bars of variable intensity. The positions of the centromeres are shown as thicker *blue* bars.

Telomeric repeats were identified at both terminal ends of four chromosomes (Chr 1, Chr 5, Chr 8, and Chr 11), at one end of ten chromosomes (Chr 3, Chr 7, Chr 9, Chr 10, Chr 13 to Chr 16, Chr 18, and Chr X, including five of the six telocentric chromosomes), and at neither end of five chromosomes (Chr 2, Chr 4, Chr 6, Chr 12, and Chr 17), indicating likely truncated assemblies at the ends of some of the chromosomes. Internal telomeric repeats containing >90 hexamers were also identified on Chr 3, Chr 6, Chr 7, Chr 9, and Chr 11 (Figure 3), and are likely the remnants of ancestral chromosomal fusion events (Thomsen et al. 1996; Kumar et al. 2017). All except one of these internal repeats is contiguously assembled; having 6,521 assembled hexameric repeats, the region on Chr 6 is the largest internal telomeric repeat in the genome and is associated with a break in assembly contiguity.

### Quantifying homozygosity

As the high contiguity of the Babraham assembly may in part be due to the homozygosity resulting from extensive inbreeding, we also sought to assess the amount of heterozygosity across the Babraham genome. A total of 671,716 single nucleotide polymorphisms (SNPs, 0.030 %) were heterozygous across the autosomes within the Babraham (P18-11073) Illumina sequencing reads (coverage depth: ∼51x) (Figure 4; Table 2). To compare between different Babraham individuals, genomic sequencing reads from the archived primary fibroblast cells of another male Babraham revealed 1,094,207 autosomal SNPs (0.048%) (coverage depth: ∼28x). These values are in contrast to the Duroc individual used to generate Sscrofa11.1 (Warr et al. 2020) in which 4,181,036 autosomal SNPs (0.185 %) were identified in that individual (coverage depth: ∼46x). Likewise, MARC1423004, the individual used to generate the USMARCv1.0 assembly was found to possess 4,121,063 autosomal SNPs (coverage depth: ∼220x). Thus, as expected given its history, the individual used to generate the Babraham pig genome is considerably more homozygous than either of the individuals used to generate the reference or the USMARCv1.0 assemblies.

**Figure 4.**
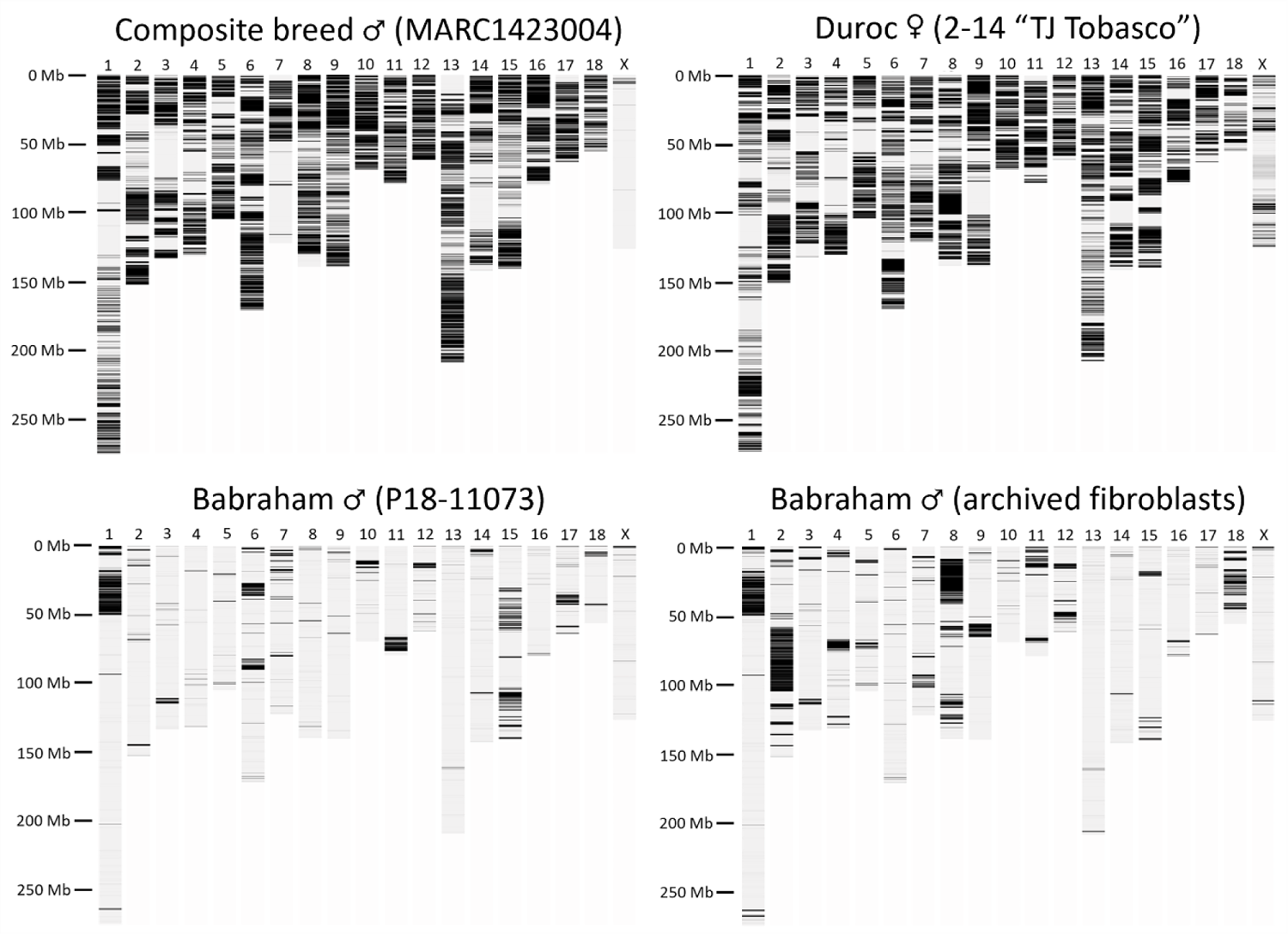
Heterozygosity of the individuals used to generate the USMARCv1.0 assembly (MARC1423004, *upper lef*t), the Sscrofa11.1 assembly (Duroc 2-14 “TJ Tobasco”, *upper right*), and the TPI_Babraham_pig_v1 assembly (P18-11073, lower left). The heterozygosity of a second Babraham individual is also shown (*lower right*) using whole genome sequencing reads generated from archival primary fibroblast cells. Reads from all individuals were mapped to Sscrofa11.1 and the number of heterozygous positions were summed and visualized using 200 kb sliding windows.

**Table 2.** Heterozygosity of animals used in pig genome assembles.

| Chr | Number of heterozygous autosomal SNPs |  |  |  |
| --- | --- | --- | --- | --- |
|  | Babraham<br>P18-11073 | Babraham<br>fibroblasts | Duroc 2-14<br>TJ Tobasco | MARC<br>1423004 |
| 1 | 149924 | 142024 | 334023 | 360418 |
| 2 | 27999 | 198839 | 291391 | 271310 |
| 3 | 21624 | 38203 | 193026 | 208336 |
| 4 | 7347 | 59090 | 177950 | 207559 |
| 5 | 12018 | 31223 | 233828 | 198982 |
| 6 | 78645 | 26059 | 295567 | 340965 |
| 7 | 47860 | 43725 | 221471 | 125536 |
| 8 | 12361 | 217931 | 295207 | 302047 |
| 9 | 14057 | 44393 | 307968 | 295318 |
| 10 | 35812 | 14295 | 193232 | 218003 |
| 11 | 52966 | 64659 | 188629 | 169332 |
| 12 | 24031 | 48358 | 140636 | 171210 |
| 13 | 11038 | 16926 | 348306 | 350704 |
| 14 | 32986 | 19045 | 264190 | 192907 |
| 15 | 91850 | 32739 | 278352 | 247408 |
| 16 | 5395 | 15828 | 178580 | 215699 |
| 17 | 33721 | 8818 | 150041 | 146865 |
| 18 | 12082 | 72052 | 88639 | 98464 |
| Total | 671716 | 1094207 | 4181036 | 4121063 |

As a measure of autozygosity (i.e., identity by descent), runs of homozygosity longer than 1 Mb (ROH_1mb_) were identified by mapping the Babraham and Duroc Illumina reads to Sscrofa11.1. To determine an allowable SNP density to include in the ROH, a background error rate was calculated using the Babraham Chr X. Except for mapping and sequencing errors, Chr X from the male Babrahams should have few or no SNPs outside the pseudoautosomal region (PAR), which comprises the first approximately 6.9 Mb (Skinner et al. 2013). Outside this PAR, the mean error rate was calculated using 200 kb windows as 1 SNP in 20 kb from the Babraham Illumina data. For both the P18-11073 and the archived fibroblast sample, this error rate varied slightly across windows, such that an upper 95^th^ percentile error rate was calculated as being approximately 1 SNP in 5 kb. Using the lower threshold of 1 SNP in 20 kb, expressed as a proportion, the ROH_1mb_ was calculated to be 0.47 (P18-11073) and 0.60 (archived fibroblasts) of the Babraham Chr X outside of the PAR. However, the upper threshold of 1 SNP in 5 kb resulted in a more expected ROH_1mb_ of 0.94 for both individuals.

Therefore, this higher error rate threshold was used to calculate the ROH_1mb_ segments across the autosomes. A total of 337 (P18-11073) and 325 (archive) ROH_1mb_ segments were identified across the Babraham autosomes, amounting to approximately 1,971 Mb (87 % of autosomal sequence) and 1,836 Mb (81 %), respectively (Table 3). In contrast, 189 ROH_1mb_ segments were identified in the Duroc autosomes totaling approximately 643 Mb, or 28 % of the autosomal sequence, and for MARC1423004 the autosomes contained 155 ROH_1mb_ segments comprising approximately 554 Mb (22 %). Thus, the Babraham pig displays a considerable amount of autozygosity due to intense inbreeding.

**Table 3.** Runs of homozygosity >1 Mb in Babraham, Duroc, and MARC individuals.

| Chr | No. of ROH segments |  |  |  | size of ROH (Mb) |  |  |  |
| --- | --- | --- | --- | --- | --- | --- | --- | --- |
|  | Babraham<br>(P18-11073) | Babraham<br>(fibroblasts) | Duroc 2-14<br>TJ Tobasco | MARC | Babraham<br>(P18-11073) | Babraham<br>(fibroblasts) | Duroc 2-14<br>TJ Tobasco | MARC<br>1423004 |
| 1 | 30 | 34 | 25 | 20 | 226 | 223 | 101 | 94 |
| 2 | 19 | 20 | 14 | 16 | 136 | 80 | 44 | 48 |
| 3 | 18 | 16 | 12 | 13 | 122 | 116 | 53 | 56 |
| 4 | 13 | 17 | 10 | 12 | 124 | 107 | 59 | 40 |
| 5 | 16 | 19 | 7 | 7 | 96 | 79 | 33 | 21 |
| 6 | 27 | 24 | 11 | 13 | 141 | 158 | 35 | 31 |
| 7 | 25 | 19 | 11 | 7 | 100 | 101 | 35 | 78 |
| 8 | 23 | 23 | 12 | 7 | 127 | 75 | 29 | 19 |
| 9 | 19 | 12 | 9 | 8 | 132 | 122 | 44 | 11 |
| 10 | 14 | 15 | 4 | 2 | 62 | 63 | 7 | 3 |
| 11 | 10 | 7 | 6 | 3 | 65 | 59 | 17 | 4 |
| 12 | 15 | 11 | 4 | 1 | 53 | 46 | 9 | 2 |
| 13 | 28 | 32 | 17 | 15 | 199 | 196 | 42 | 39 |
| 14 | 24 | 26 | 12 | 7 | 130 | 134 | 40 | 55 |
| 15 | 23 | 19 | 11 | 11 | 90 | 125 | 33 | 24 |
| 16 | 11 | 11 | 9 | 5 | 72 | 70 | 23 | 14 |
| 17 | 15 | 13 | 7 | 4 | 47 | 57 | 18 | 7 |
| 18 | 7 | 7 | 9 | 4 | 49 | 26 | 21 | 7 |
| X | 20 | 24 | 14 | 7 | 112 | 116 | 38 | 120 |
| 1 to 18 | 337 | 325 | 190 | 155 | 1971 | 1837 | 643 | 554 |
| Total | 357 | 349 | 204 | 162 | 2083 | 1952 | 680 | 674 |

### Immune-related gene complexes are largely contiguous

Due to their repetitive nature, immune-related gene complexes are often poorly assembled in whole genome sequencing efforts. The nature of somatically rearranging B cell and T cell receptor genes also potentially complicates genome assemblies across these regions when using genomic DNA derived from blood. To mitigate this, we selected the largely immune-privileged cerebral cortex as a source of genomic material for the present study. Given the utility of the inbred Babraham pig for immunological studies, we sought to examine several immune-related genomic regions that are functionally important in lymphocyte immunobiology and commonly misassembled in whole genome sequencing efforts.

### The T cell receptor (TCR) loci

The pig TCR alpha and TCR delta chains are encoded within the same gene cluster, TRA/D (Babraham Chr 7: 76,710,039 – 77,541,877). This is the largest and most gene-dense region presently described, spanning approximately 1 Mb and containing approximately 118 *TRAV* and *TRDV* gene segments, and is rarely contiguously assembled. In the Babraham assembly there are two sequence gaps and one 96 kb unplaced contig (contig_21). In Sscrofa11.1 (position: 7: 76,471,214 – 77,539,127) there is one sequence gap and a 43 kb unplaced contig (GenBank accession: AEMK02000555). Specific details regarding individual genes and polymorphisms are complicated by disruptions in the assemblies and the high similarity between many of the V gene segments. A ∼73 kb duplication within the V region is present in Sscrofa11.1, but not the Babraham; and another ∼95 kb duplication is found in both (Figure 5). Peculiarly, these duplicated regions are not in the same locations in both assemblies. However, the actual organization is difficult to determine as the sequence gaps in both assemblies are all adjacent to these duplications (Figure 5), thus implicating these duplications and their repetitiveness to the lack of contiguity across the V region.

**Figure 5.**
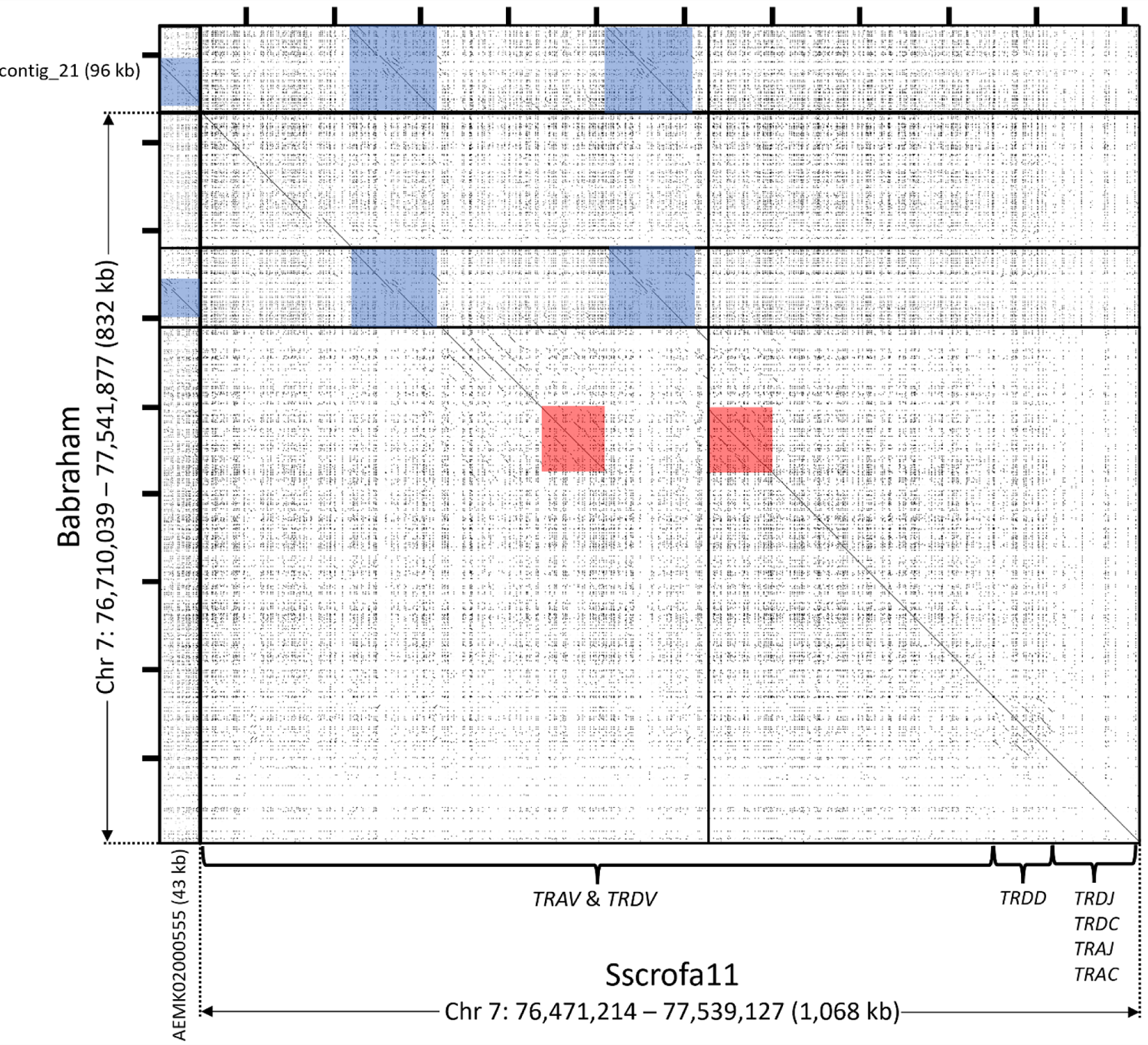
Recurrance plot comparison of the TRA/D locus in the Babraham (*vertical axis*) and Sscrofa11.1 (*horizontal axis*) assemblies. Gaps in the Babraham and Sscrofa assemblies are indicated by thick horizontal and vertical lines, respectively. Unplaced contigs in both assemblies are depicted here upstream from the V region. A ∼73 kb region that is duplicated in the Sscrofa11.1 assembly, but not the Babraham, is shaded *red*; and a ∼95 kb region that is duplicated in the Babraham assembly and triplicated in Sscrofa11.1 is shaded *blue*. Tick marks on top and at left are each separated by 100 kb.

The TCR beta chain (TRB) region (Babraham Chr 18: 7,345,551 – 7,686,331) has been previously described for the Sscrofa11.1 assembly (Chr 18: 7,397,804 – 7,734,192) (Massari et al. 2018). Within that assembly, the *TRB* is intact on a single contig that spans the entire chromosome (∼56 Mb), whereas a single sequence gap disrupts the *TRB* in the Babraham assembly – the only such sequence gap in the Chr 18 assembly. The Sscrofa11.1 *TRB* region contains 38 described *TRBV* genes (Massari et al. 2018) compared to 36 *TRBV* genes in the Babraham assembly. Recurrence plot analysis comparing the two assemblies revealed two distinct *TRBV* regions containing highly repetitive sequence (Figure 6). Of these, the *TRBC*-distal region is variable in gene content containing ten *TRBV* genes in Sscrofa11.1 (*TRBV4-1* to *TRBV2-5*), but only eight in the Babraham. The *TRBC*-proximal region contains three highly similar *TRBV* genes (*TRBV20-1* to *TRBV20-3*) in both assemblies, plus an L1 insertion in the Babraham. This C-proximal cluster also abuts the Babraham sequence gap and thus the sequence similarity within this gene cluster presumably contributed to the disruption of the Chr 18 assembly.

**Figure 6.**
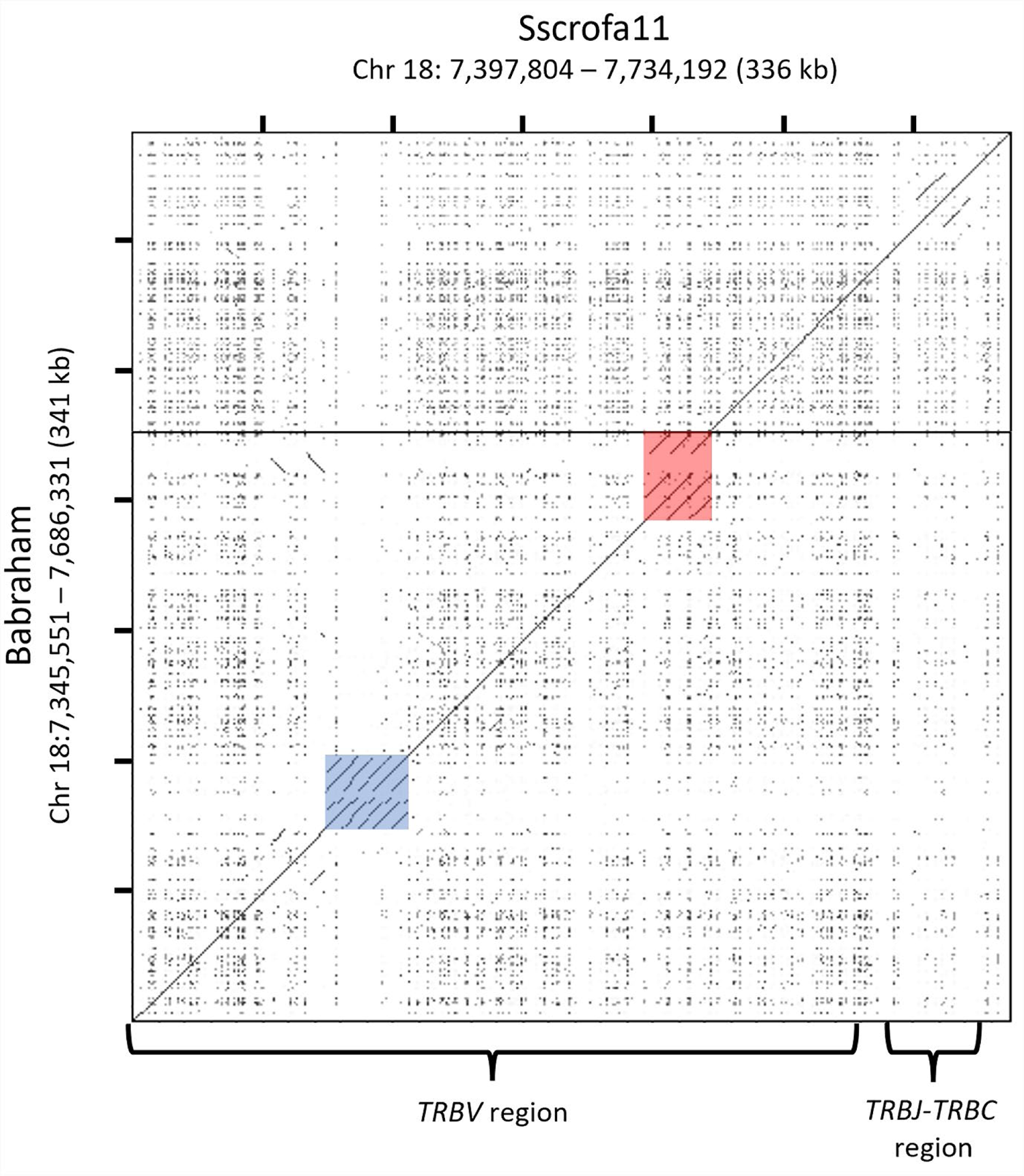
Recurrance plot comparion of the TRB locus in the Babraham (*vertical axis*) and Sscrofa11.1 (*horizontal axis*) assemblies. A single sequence gap in the Babraham assembly – the only such gap on Chr 18 – is indicated as a thick horizontal line. This sequence gap is adjacent to a ∼26 kb (Sscrofa11) to ∼34 kb (Babraham) region containing three tandemly duplicated *TRBV* paralogs present in both assemblies (region shaded in *red*). In the Babraham, this region is larger due to an additional L1 insertion. Another ∼32 kb region (shaded in *blue*) containing 10 closely related *TRBV* paralogs in Sscrofa11.1 appears to vary in gene content between haplotypes, as the same region only contains eight *TRBV* genes in the Babraham assembly.

The pig TCR gamma chain (TRG) region (Babraham Chr 9: 108,295,979 – 108,409,334) has recently been described in detail for the Babraham, Sscrofa11.1, and USMARCv1.0 assemblies (Le Page et al. 2021; Linguiti et al. 2022). In the Babraham, this region is intact and in the middle of a 41.7 Mb contig. The region contains four polymorphic V-J-C gene cassettes in both the Babraham and Sscrofa11.1 (Chr 9: 108,678,980 – 108,791,795) assemblies, although only three cassettes were identified in the USMARCv1.0 assembly (Chr 9: 30,653,846 – 30,739,227) (Le Page et al. 2021). Although the first of these cassettes was found to be the most abundantly expressed in general, *TRGV6* (of the second cassette) was previously found to be the single-most transcribed V gene segment; and while *TRGV6* is functional in the Babraham, it is putatively non-functional in the other porcine assemblies, due to being out-of-frame (Le Page et al. 2021).

### The B cell receptor (BCR) loci

The immunoglobulin heavy chain (IGH) region (Babraham Chr 7: 125,292,945 – 125,642,007) is assembled to the telomeric end of Chr 7 on a 46 Mb contig, confirming previous cytogenetic evidence for its localization (Yerle et al. 1997). This region is unplaced in previous pig reference assemblies. Within Sscrofa11, the *IGH* region is split across at least six unplaced contigs (GenBank: AEMK02000149, AEMK02000151, AEMK02000188, AEMK02000452, AEMK02000566, and AEMK02000599); in particular, the Sscrofa11.1 IGH constant region and four *IGHV* genes are assembled to the end of a 3.8 Mb contig (GenBank: AEMK02000452). In pigs, this region is variable in *IGHG* content (and thus IgG isotypes). *IGHG1*, *IGHG3*, and *IGHG4* seem to be found in all haplotypes, whereas six additional *IGHG* genes have been found to be variably present depending on the haplotype (Zhang et al. 2020). The Babraham assembly itself contains *IGHG1*, *IGHG3*, and *IGHG4*, as well as *IGHG2a*, which is a close paralog of *IGHG4*. In contrast, the unplaced contiguous Sscrofa11.1 sequence contains the same four *IGHG* as the Babraham, in addition to *IGHG5a* and *IGHG2c*. A total of 25 *IGHV* gene segments, including 13 that are putatively functional, are present in the Babraham assembly. The *IGHV* gene most distal to the constant region sits a mere 4 kb from the telomeric-end of the assembly, and since the flanking telomere is not present, the assembled *IGHV* region is possibly incomplete. A BLAST survey identified three additional small unplaced contigs (contig_547, 1.5 kb; contig_1142, 7.6 kb; and contig_1640, 29.1 kb) containing one, one, and four *IGHV* pseudogenes, respectively. These may represent either additional constant region-distal gene segments or alternative alleles that could not be assembled.

The immunoglobulin lambda light chain (IGL) region (Babraham Chr 14: 48,527,945 – 48,766,643) is continuous within a 15.2 Mb contig and falls within a 16 Mb ROH in both Babraham Illumina datasets. This region was previously characterized using overlapping bacterial artificial chromosomes (BACs) derived from the same Duroc individual used to generate the reference assembly, Sscrofa11.1 (Schwartz et al. 2012b). The IGL region is known to be polymorphic and possibly variable in gene content, as evidenced by *IGLV3-6* which can be present as either a null allele or as a highly transcribed functional allele (Schwartz and Murtaugh 2014; Guo et al. 2016). This diversity is apparent in the Babraham as well since both *IGLV3-6* and the adjacent *IGLV3-2* are deleted. The *IGLC* region likewise appears to be variable in gene content. The previous BAC characterization revealed three *IGLJ*-*IGLC* cassettes and *IGLJ4* with no corresponding downstream *IGLC* (Schwartz et al. 2012b). The IGL region within the Sscrofa11.1 assembly (Chr 14: 48,741,433 – 49,012,235), however, contains four intact cassettes, plus *IGLJ4*, and peculiarly the Babraham assembly contains six *IGLJ*-*IGLC* cassettes, as well as *IGLJ4*. In all assemblies, the most 5’ *IGLJ* contains the same non-canonical “FSGS” motif as described for *IGLJ1*, and the remaining cassettes all possess the same 1.3 kb spacing and canonical “FGGG” motif as described for the *IGLJ2* and *IGLJ3* gene segments, indicating that the more distal 3’ *IGLJ*-*IGLC* cassettes with canonical *IGLJ* are particularly prone to expansion and/or contraction.

The immunoglobulin kappa light chain (IGK) region (Babraham Chr 3: 57,436,231 – 57,625,777) is fragmented by two sequence gaps within the repetitive *IGKV* region. This includes a small (11.9 kb) intervening contig flanked by two much larger contigs containing the 5’ and 3’ ends of the region. In contrast the same region in Sscrofa11.1 (Chr 3: 57,118,524 – 57,321,145) is continuous. As with the IGL, this region was previously characterized using BAC sequences derived from the same Duroc individual used to generate Sscrofa11.1 (Schwartz et al. 2012a). However, that characterization was incomplete, as it only identified the 14-most *IGKC*-proximal *IGKV* gene segments. We have therefore characterized the IGK gene content in both the Babraham and Sscrofa11.1 assemblies, the latter of which is continuous, and identified 23 *IGKV* gene segments in Sscrofa11.1 and 19 *IGKV* in the Babraham assembly, although at least two of these, *IGKV2-13* and *IGKV1-14* may be missing in a sequence gap, a BLAST search of unplaced contigs did not identify them.

### The Leukocyte Receptor Complex (LRC)

The LRC (Babraham Chr 6: 58,236,196 – 58,935,786) is continuous in the Babraham assembly, but disrupted in Sscrofa11.1 (Chr 6: 55,898,983 – 59,234,370) by the presence of a sequence gap and large inversion due to mis-assembly within a 197 kb sub-region that contains 17 repetitive leukocyte immunoglobulin-like receptor (*LILR*) genes and fragments from two distinct sub-families (Schwartz and Hammond 2018). In contrast, the Babraham assembly contains fewer *LILR* than Sscrofa11, with only 11 genes, including two gene fragments. Compared to our previous characterization of the LRC in Sscrofa11.1, the identified genes in the Babraham correspond to *LILR1B1* and *LILR2B8* to *LILR1A16*, with *LILR2B2* to *LILR1A7* being absent from the Babraham genome. Despite this, all six putatively functional genes in the Sscrofa11.1 assembly are also functional in the Babraham; and in addition to these, *LILR2B8*, which is putatively non-functional in Sscrofa11.1, is putatively functional in the Babraham. The remaining genes of the LRC, including the gene content variable novel immunoglobulin-like receptor genes, are similar to the described Sscrofa11.1 assembly (Schwartz and Hammond 2018).

### The Natural Killer Complex (NKC)

The Babraham NKC (Babraham Chr 5: 63,923,511 – 65,716,322) is continuous within a 23.4 Mb contig and within a >5 Mb ROH in both Babraham Illumina datasets. This region is likewise contiguous within Sscrofa11.1 (Chr 5: 61,441,125 – 63,228,372). Although highly expanded in bovids, equids, and rodents, the killer cell C-type lectin-like receptor (KLR) genes appear to represent a minimal set of genes in the pig, including only one inhibitory *KLRC* gene which is otherwise expanded in all studied species, including humans which have four *KLRC* genes (Schwartz et al. 2017). Furthermore, we found no indication of gene content variation across this region between the two assemblies.

### The Major Histocompatibility Complex (MHC)

The MHC class I (Babraham Chr 7: 23,090,615 – 23,868,138) and class II (Babraham Chr 7: 25,057,296 – 25,415,322) regions are separated by the MHC class III region which also includes the centromere and two associated sequence gaps which may contain additional unplaced sequence. We previously determined that Babraham pigs are homozygous for the MHC haplotype Hp-55.6 (Schwartz et al. 2018), which is confirmed in the present assembly. In addition to the previously described alleles for *SLA-1*, *SLA-2*, *SLA-3*, *SLA-6*, *SLA-7*, and *SLA-8* within the extended MHC class I region, we further identified additional pseudogenes for *SLA-1*, *SLA-4*, and *SLA-5*, as well as functional *SLA-11*. Moreover, *SLA-6* was found to possess a deletion encompassing all of exon 1, with no potential alternative leader exon identified. The designation of Babraham *SLA-6* as a null allele is consistent with our earlier finding that all cDNA sequences for *SLA-6* were unspliced (Schwartz et al. 2018).

The MHC class II region is found on the long-arm of Chr 7 approximately 180 kb from the centromere which bisects the class III region. In addition to the described class II alleles for *SLA-DRB1* and *SLA-DQA* within the Hp-55.6 haplotype, we resolved the allele designation for *SLA-DQB* as being *SLA-DQB*08:01* and additionally identify the *SLA-DRA* allele as *SLA-DRA*02:02:03*. *SLA-DRB4*, although classed as a pseudogene and currently not represented within the ImmunoPolymorphism Database (IPD)-MHC (Maccari et al. 2020), is putatively functional in the both the Babraham and Sscrofa11.1 assemblies, although future work is necessary to determine whether it is functionally transcribed and translated.

## Discussion

At a cost of tens of thousands of US dollars, the presently described PacBio long-read Babraham pig assembly, error-corrected with Illumina short reads, is more contiguous (contig N50 = 34.9 Mb) than the initial Sscrofa11 PacBio assembly (contig N50 = 14.5 Mb) that was generated prior to gap filling which included the earlier sequencing data and Nanopore reads, and slightly less than the final Sscrofa11.1 assembly (Warr et al. 2020).

Divergent haplotypes can negatively affect an assembly’s contiguity due to their competition for assembly into a haploid representation of a diploid genome. Thus, homozygosity should aid whole genome assembly, and recent approaches have therefore sought to limit the effect that heterozygosity has on contiguity. This includes using individuals from genetically isolated and/or bottlenecked populations (Bickhart et al. 2017), or by generating two distinct haploid assemblies from an offspring with genetically divergent parents (Koren et al. 2018; Low et al. 2020; Rice et al. 2020; Bredemeyer et al. 2021). It is therefore plausible that the extreme homozygosity of the sequenced Babraham individual contributed to the relatively high contiguity of the currently described assembly.

Advancements over the last decade in long-read sequencing technologies and improved scaffolding techniques have allowed for dramatic improvements in the contiguity of whole genome assemblies at a greatly reduced economic cost. The completion of the pig reference genome, Sscrofa9, in 2009 was the result of an extensive global effort which used 4x to 6x Sanger whole genome shotgun (WGS) reads mostly derived from the CHORI-242 BAC library (Archibald et al. 2010) and achieved a contig N50 of 54.2 kb with extensive manual finishing and gap filling. The reference was later updated to Sscrofa10.2 (contig N50 = 576 kb) with >30x Illumina GAII short-read WGS mostly based on CHORI-242 (Groenen et al. 2012), and recently updated to Sscrofa11.1 (contig N50 = 48.2 Mb) with 65x WGS PacBio RSII reads, error-corrected with Illumina HiSeq 2500 WGS reads, and gap filled using both Oxford Nanopore and Sanger reads derived from CHORI-242 (Warr et al. 2020). Chromosome assignment of Sscrofa11.1 (and USMARCv1.0) scaffolds, which we also based the Babraham chromosomal assignments on, was itself initially based on the earlier Sscrofa10.2 assembly (Groenen et al. 2012), and ultimately on earlier physical mapping data (Humphray et al. 2007). Thus, any scaffolding errors present in the earlier reference assemblies, including contig ordering and orientation, would have carried through to the current pig genome assemblies, including for the Babraham.

Chr Y is highly repetitive and predicted to be approximately 30 Mb in the pig (Skinner et al. 2016). As a result of the repetitiveness and difficulty in assembling it, Chr Y is often excluded from mammalian genome assemblies. In the Babraham assembly, Chr Y is incompletely assembled to a 5.5 Mb scaffold that is poorly contiguous compared to the rest of the Babraham assembly. Therefore, much of the Chr Y sequence is expected to be represented amongst the unplaced contigs. Despite this, there is less unplaced sequence overall in the Babraham assembly (∼40 Mb) compared to either Sscrofa11.1 (∼65 Mb) or USMARCv1.0 (∼330 Mb). Since much of this unplaced sequence is also expected to derive from alternative haplotypes (Koren et al. 2018), the relatively low amount of unplaced sequence likely reflects the high homozygosity of the sequenced Babraham pig.

In 1999, restriction fragment fingerprinting suggested a similar level of homozygosity in the Babraham pig as inbred mice (Signer et al. 1999), and after multiple generations of continued inbreeding, extensive genome-wide homozygosity was further confirmed in 2016 from the SNP genotyping of five Babraham individuals (Nicholls et al. 2016). The extent of homozygosity and the remaining regions of heterozygosity identified in that study mirror our present findings using whole genome short-read data; in particular, relatively extensive tracts of heterozygosity remain in some, but not all, Babraham individuals on Chr 2 and Chr 8. Such genetic variation may contribute to the phenotypic variation between Babraham individuals; however, overall phenotypic variation is greatly reduced compared to other large pig breeds.

Of the immune-related gene complexes investigated, only the IGL and NKC lie within an ROH in both the Babraham individuals that we examined. This is despite sequencing of individual MHC-I and MHC-II alleles that indicates homozygosity across those regions in all animals sequenced so far (Schwartz et al. 2018). However, because these gene complexes tend to be highly repetitive, and thus notoriously difficult to accurately assemble and map using short read data, at least some of the limited heterozygosity observed in these regions is likely the result of mis-mapping. Thus, due to inevitable short-read mis-mapping errors, our results likely underestimate homozygosity to some extent.

The *LILR* genes are the most complex of the pig LRC and have undergone recent expansions, as evidenced by the presence of many highly similar and tandemly repeated genes. It is therefore highly plausible for *LILR* gene content variation to exist between different haplotypes. This gene content variation may explain why the Babraham has fewer apparent *LILR* genes compared to the Sscrofa11.1 assembly. The homozygosity across the LRC in the sequenced Babraham may have eased the assembly across this region into a single contig, while the heterozygous Sscrofa11.1 assembly was disrupted (Schwartz and Hammond 2018).

The pig *TRA/D* locus at approximately 1 Mb is similar in scale to the human (1 Mb) and dromedary camel (877 kb) (Massari et al. 2021), but substantially less than bovines (3.5 Mb) (Connelley et al. 2014). This locus is considerably larger than any of the other somatically rearranging T cell and B cell receptor genes, and due to the large (∼73 kb and ∼95 kb) repeat structures, it remains particularly challenging to completely assemble. In contrast, the IGH locus of the Babraham assembly possibly represents the first completely assembled porcine IGH region and is correctly assembled to the telomeric end of the long arm of Chr 7 (Yerle et al. 1997). Although it remains to be verified if all Babrahams share the same IGH haplotype, the sequenced individual possesses four *IGHG* genes, including the variably present *IGHG2a*. While not found in all pigs, the expressed IgG2a subclass has recently been shown to have strong Fc binding to NK cells, and strong effector functions, including complement-dependent cellular cytotoxicity, antibody-dependent cellular phagocytosis, and degranulation of NK cells (Paudyal et al. 2022). The antibody light chain gene segments *IGLV3-2* and *IGLV3-6* are deleted in the sequenced Babraham haplotype, and similar variation was previously shown to skew the expressed IGL repertoire in favor of different gene segments (Schwartz 2013; Schwartz and Murtaugh 2014; Guo et al. 2016).

Of the immune-related gene complexes that we examined, only the non-classical MHC genes and the NKC region appear to be fixed in gene content between pigs. This potentially extensive haplotypic variation across these regions could thus have profound effects on the expressed porcine immunome and variable immune phenotypes between individuals. Due to this genomic variability, the utility and availability of genomic resources matched to an experimental animal model, such as the Babraham pig, is worth considering during experimental design.

The presently described genome assembly represents an alternative porcine genome assembly of a highly inbred Babraham pig based on the Large White commercial breed. Likely due to the high amount of homozygosity resulting from inbreeding, the assembly is nearly as contiguous as the current pig reference assembly, Sscrofa11.1. Additionally, several immune-related gene complexes are more intact, making it a valuable alternative genomic resource for porcine research. The Babraham pig itself is a proven biomedical and veterinary model, due to both its high level of inbreeding and lack of variation in the MHC and likely other immunogenetic loci. The genome assembly is therefore expected to further aid research in which controlling for this genetic variability is of paramount importance.

## Methods

### Animal use and ethics statement

A representative adult male Babraham pig (animal ID: P18-11073), whose parents were half siblings, was culled from The Pirbright Institute’s herd held at the Animal and Plant Health Agency (APHA; Addlestone, United Kingdom) as part of routine herd maintenance. This procedure was carried out in accordance with the UK Animal (Scientific Procedures) Act 1986 and approved by both The Pirbright Institute Animal Welfare and Ethical Review Body and the APHA Animal Welfare and Ethics Committee.

### Nucleic acid purification and sequencing

Tissue from the frontal lobe of the cerebral cortex was chosen for whole genome sequencing due to its inherent lack of immune cells with rearranging receptors (i.e., B cells and T cells), which may complicate assembly efforts across these respective genetic loci. A sample of the tissue was transported to the University of Utah Core Research Facilities (Salt Lake City, Utah) on dry ice for high molecular weight genomic DNA purification and sequencing. For genome assembly, long-read sequencing was performed using the PacBio Sequel II platform which resulted in 11,141,834 reads with an average read length of 12,552 bp (∼57x coverage). For error correction, short-read sequences were generated using the Illumina TruSeq DNA PCR-Free library preparation kit and the Illumina NovaSeq 6000 platform which resulted in 415,666,795 paired-end 150 bp reads (∼51x coverage).

Additional genomic DNA was prepared from primary fibroblast cells collected from a male Babraham pig and archived at The Pirbright Institute circa 2015. Approximately 3 x 10^7^ cells were resuspended in 5 ml of PBS and lysed with 25 ml lysis buffer (140 mM NH_4_Cl and 17 mM TRIS-HCl, pH 7.4). The resulting pellet was then resuspended in 9 ml (10 mM TRIS-HCl, 400 mM NaCl, 2 mM EDTA, pH 8.0) and digested for one hour at 37 °C after the addition of 10 % sodium dodecyl sulfate (600 ul) and 100 mg/ml RNase A (13 ul). Nucleases were then inactivated with the addition of 20 mg/ml Proteinase K (100 ul) for eight hours. High molecular weight genomic DNA was then precipitated by adding 6 M NaCl (3 ml), centrifuging, treating the supernatant with two volumes (∼26 ml) of 100 % ethanol, and centrifuging again to produce a DNA pellet that was further purified using 80 % ethanol. The final pellet was resuspended in 0.1x TE buffer (1 mM TRIS and 0.1 mM EDTA, pH 8.0) and quantified using an Agilent TapeStation 4150. As above, DNA was provided to the University of Utah Core Research Facilities for Illumina TruSeq DNA PCR-Free library preparation and sequencing using a HiSeq 2500 which generated 278,898,802 paired-end 125 bp reads (approximately 28x coverage).

### Genome assembly and error correction

The PacBio Sequel II sequencing reads were *de novo* assembled into contigs and scaffolded using Flye, v2.5 (Kolmogorov et al. 2019) with parameters set to: --asm-coverage 30 -t 30 and error-corrected using Pilon (version 1.24) (Walker et al. 2014) and the P18-11073 Illumina sequences. The error-corrected contigs/scaffolds were then mapped to the Sscrofa11.1 chromosomal assembly (GenBank: GCA_000003025.6) using Minimap2 (Li 2018). This mapping was used to order and orient the Babraham contigs into chromosomes, in which the *de novo* assembled contigs and scaffolds were separated by a span of 100 N’s. Orientation and identity were confirmed by mapping these chromosomal assemblies back to Sscrofa11.1 using Minimap2 with the preset parameter -x asm5 for long assembly to reference mapping with up to 5 % sequence divergence (Li 2018). The Minimap2 output in pairwise mapping format (paf) was then visualized for each chromosome in R (v3.4.1) using dotPlotly with parameters set to: -m 100 -q 50000 (Poorten). The 1,035 unplaced contigs were screened for contaminating sequence using Kraken (version 1.1.1) and the complete Kraken database including viral, bacterial, and fungal sequence (Wood and Salzberg 2014). This flagged 378 contigs as potentially containing contaminating viral or bacterial sequence. However, all except two of these successfully mapped to Sscrofa11.1 using Minimap2, indicating that the Kraken hits were false positives. The remaining two unmapped contigs fully contained relatively simple (i.e., A(C_n_) _n_ and (TTTAAC) _n_) repeats. Thus, all 1,034 unplaced contigs were retained in the final assembly.

### Analysis of heterozygosity

Short-read whole genome sequencing reads were mapped to Sscrofa11.1 using the Burrows-Wheeler Aligner (BWA; version 0.7.12) (Li and Durbin 2009). For the Babrahams, this included both the 4.16 x 10^8^ reads from P18-11073 and the 2.79 x 10^8^ reads from the primary Babraham fibroblast cells described above. For the Duroc (i.e., “TJ Tobasco”), FASTQ files collectively containing approximately 3.74 x 10^8^ Illumina HiSeq 150 bp paired-end sequencing reads (∼46x coverage) were acquired from BioProject accession PRJEB9115. Sequences for MARC1423004, the individual used to generate the USMARCv1.0 assembly, were acquired from the sixteen NextSeq 500 runs archived within BioProject accession PRJNA392765 and totaled 1.79 x 10^9^ paired-end 150 bp sequencing reads (∼220x coverage). Variant sites were identified using SAMtools (version 1.2) and BCFtools (version 1.3.1) (Li et al. 2009; Li 2011a), and the resulting VCF files were indexed with Tabix (version 1.10.2-45-gb22e03d) (Li 2011b). Only those SNPs with a Phred-scaled QUAL score ≥30 were considered for further analyses. For the Babraham and MARC1423004 sequences, the total number of bi-allelic (ALT/REF) and tri-allelic (ALT1/ALT2) heterozygous SNPs were summed within each 200 kb window. For the Duroc, any tri-allelic SNPs would be the result of mapping error, so only the total number of heterozygous bi-allelic (ALT/REF) SNPs were summed for each 200 kb window. Heterozygosity across the genome was then visualized using Gitools version 2.3.1 (Perez-Llamas and Lopez-Bigas 2011).

### Telomeric and centromeric repeats

Telomeric repeats were identified by searching for repeat sequences containing exact matches of at least three tandem hexamers of either TTAGGG or CCCTAA using Tandem Repeats Finder (TRF) version 4.09 (Benson 1999). The number of hexamers within each identified repeat was summed and visualized across each chromosome using Gitools version 2.3.1 (Perez-Llamas and Lopez-Bigas 2011) and a window size of 200 kb. Output from TRF was also used to identify large (i.e., 10 kb (chr15) to 552 kb (chr2)) centromeric repeat regions in the expected chromosomal locations based on previous analyses (Hansen 1977; Warr et al. 2020). Gitools was also used to visualize these centromeric repeat regions within each chromosomal assembly.

### Annotation of immune-related gene complexes

Assembled chromosomes and unplaced contigs were queried using both the basic local alignment search tool (BLAST) (Altschul et al. 1990) for genes of interest within the natural killer complex (NKC), leukocyte receptor complex (LRC), major histocompatibility complex (MHC), and T cell and B cell receptor loci using previously reported characterizations or from IPD-MHC (Lunney et al. 2009; Eguchi-Ogawa et al. 2012; Schwartz et al. 2012a; Schwartz et al. 2012b; Maccari et al. 2017; Schwartz et al. 2017; Massari et al. 2018; Schwartz and Hammond 2018; Schwartz et al. 2018; Hammer et al. 2020; Maccari et al. 2020; Le Page et al. 2021), which the Babraham was compared to. This was aided with the use of the conserved domain search tool (Marchler-Bauer and Bryant 2004; Marchler-Bauer et al. 2015) to help identify additional genes and gene fragments. Exons were manually annotated within the chromosomal assemblies using Artemis (version 17.0.1) (Rutherford et al. 2000). MHC alleles were named based on their identity to known alleles within IPD-MHC (Maccari et al. 2017; Maccari et al. 2020). Recurrence plot comparisons of gene loci between the Babraham and Sscrofa11.1 assemblies were generated using Dotter (version 4.44.1) (Sonnhammer and Durbin 1995).

## Data Access

The TPI_Babraham_pig_v1 genome assembly is available from ENA/GenBank under the accession GCA_031225015.1. Illumina and PacBio data used to generate the genome assembly have been submitted to the NCBI BioProject database (https://www.ncbi.nlm.nih.gov/bioproject/) under accession number PRJNA1009406. Illumina data generated from the archived Babraham primary fibroblast cells have also been submitted to the NCBI BioProject database under accession number PRJNA992241. Specific allele sequences described in the text and manually annotated for the immune-related gene complexes in the Babraham assembly are available from the authors upon request. Babraham pigs are a UK national capability resource managed by The Pirbright Institute (Woking, UK). Individuals or groups seeking access to the Babraham pig herd are encouraged to contact.

## Competing Interest Statement

The authors declare no competing interests.

## Acknowledgements

We thank Dr. Ryan Waters (The Pirbright Institute) for his helpful comments and for coordinating the collection of animal tissues with the veterinary services team at APHA. We also thank Dr. Liz Reid (formerly at The Pirbright Institute) for providing the primary Babraham fibroblast cells and Dr. Martina Hadrovic (formerly at The Pirbright Institute) for technical assistance. We thank Derek Warner from the University of Utah Core Sequencing Facility for all his effort in management of sequence acquisition. JCS, GF, and JAH are supported by United Kingdom Research and Innovation Biotechnology and Biological Sciences Research Council (UKRI-BBSRC) awards BBS/E/I/00007031, BBS/E/I/00007038, BBS/E/I/00007039, and BB/S506680/1. AKS is a Wellcome Investigator (220295/Z/20/Z).

## Author contributions

JCS, AKS, JAH, and JDP planned and coordinated the study. JDP oversaw the genomic sequencing. CPF performed the genome assembly and error correction. JCS and GF performed additional analyses. JCS wrote the manuscript which all authors read and provided input on.

## Notes

### Competing Interest Statement

The authors have declared no competing interest.

